# Ion-channel degeneracy and heterogeneities in the emergence of signature physiological characteristics of dentate gyrus granule cells

**DOI:** 10.1101/2024.02.15.580453

**Authors:** Sanjna Kumari, Rishikesh Narayanan

## Abstract

Complex systems are neither fully determined nor completely random. Biological complex systems, including single neurons, manifest intermediate regimes of randomness that recruit integration of specific combinations of functionally segregated subsystems. Such emergence of biological function provides the substrate for the expression of degeneracy, the ability of disparate combinations of subsystems to yield similar function. Here, we present evidence for the expression of degeneracy in morphologically realistic models of dentate gyrus granule cells (GC) through functional integration of disparate ion-channel combinations. We performed a 45-parameter randomized search spanning 16 active and passive ion channels, each biophysically constrained by their gating kinetics and localization profiles, to search for valid GC models. Valid models were those that satisfied 17 sub- and supra-threshold cellular-scale electrophysiological measurements from rat GCs. A vast majority (>99%) of the 15,000 random models were not electrophysiologically valid, demonstrating that arbitrarily random ion-channel combinations wouldn’t yield GC functions. The 141 valid models (0.94% of 15,000) manifested heterogeneities in and cross-dependencies across local and propagating electrophysiological measurements, which matched with their respective biological counterparts. Importantly, these valid models were widespread throughout the parametric space and manifested weak cross-dependencies across different parameters. These observations together showed that GC physiology could neither be obtained by entirely random ion-channel combinations nor is there an entirely determined single parametric combination that satisfied all constraints. The complexity, the heterogeneities in measurement and parametric spaces, and degeneracy associated with GC physiology should be rigorously accounted for, while assessing GCs and their robustness under physiological and pathological conditions.

**GRAPHICAL ABSTRACT:** 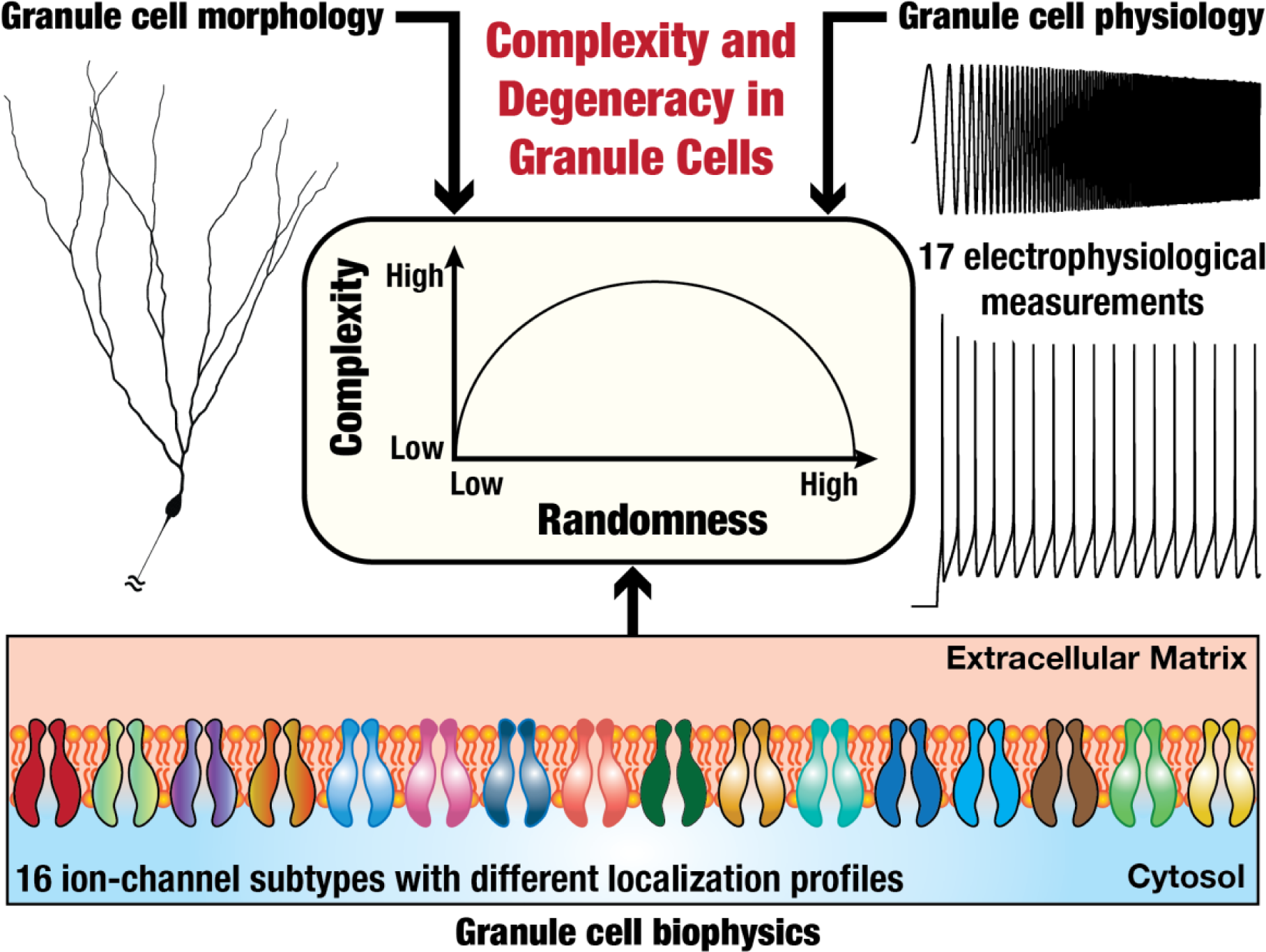

## INTRODUCTION

The dentate gyrus (DG) is the gateway to the hippocampus proper and plays critical roles in engram formation, pattern separation, and spatial navigation. The granule cells are the principal excitatory neurons in the DG that receive afferent inputs from the entorhinal cortices through the perforant pathway and send outputs to the CA3 through the mossy fibers. Electrophysiological properties of DG granule cells and their dependencies on individual ion channels manifest pronounced heterogeneities (Beining et al., 2017b; Mishra and Narayanan, 2019; 2020; Zhang and Jonas, 2020; Zhang et al., 2020; Huckleberry and Shansky, 2021; Mishra and Narayanan, 2021b; c; Schneider et al., 2023). The traditional use of a single hand-tuned model to computationally study granule cells would not accommodate such heterogeneities in and differential dependencies of characteristic physiological approaches. In addition, reliance on a single hand-tuned model would yield biased conclusions as the hand-tuned model is simply one instance of the several possible realizations of characteristic neuronal properties. The population-of-models approach is now an established route to systematically study heterogeneities and differential dependencies of functional outcomes in several biological neuronal subtypes (Foster et al., 1993; Goldman et al., 2001; Prinz et al., 2003; Marder and Taylor, 2011; Rathour and Narayanan, 2012; 2014; Basak and Narayanan, 2018; Migliore et al., 2018; Mittal and Narayanan, 2018; Rathour and Narayanan, 2019; Basak and Narayanan, 2020; Goaillard and Marder, 2021; Roy and Narayanan, 2021; Mittal and Narayanan, 2022; Arnaudon et al., 2023; Nagaraj and Narayanan, 2023; Reva et al., 2023; Roy and Narayanan, 2023), including DG granule cells (Beining et al., 2017b; Mishra and Narayanan, 2019; 2021c; Schneider et al., 2023).

The ability of disparate combinations of different ion channels to elicit similar signature physiological characteristics has been referred to as ion-channel degeneracy (Edelman and Gally, 2001; Rathour and Narayanan, 2019; Goaillard and Marder, 2021). There are several electrophysiological (Mishra and Narayanan, 2021b) and computational (Beining et al., 2017b; Mishra and Narayanan, 2019; 2021c; Schneider et al., 2023) lines of evidence for the manifestation of ion-channel degeneracy in the manifestation of signature granule cell physiology. However, the lines of computational evidence for ion-channel degeneracy in DG granule cells come either from single compartmental models or without an extensive set of electrophysiological measurements that severely constrain the physiological outcomes. Morphologically realistic models with a broad set of physiological constraints are essential because morphology sets strong structural constraints on neuronal physiology (Mainen and Sejnowski, 1996; Krichmar et al., 2002; Narayanan and Chattarji, 2010; Dhupia et al., 2015; Basak and Narayanan, 2020) and the number of physiological constraints contribute to model complexity (Yang et al., 2022; Schneider et al., 2023).

In this study, we build a population of GC models adapted from an extensive model (Beining et al., 2017b) to demonstrate ion-channel degeneracy in morphologically realistic granule cell models that were biophysically and physiologically constrained. We used a systematic and unbiased search of a 45-parameter space that spanned all ion channels and their precise subcellular distributions. Importantly, we validated each random model using 17 different sub- and supra-threshold electrophysiological measurements from DG granule cells (Mishra and Narayanan, 2020). We found a small subset (< 1%) of 15,000 randomly generated morphologically realistic models to satisfy all 17 electrophysiological constraints. We performed quantitative analyses on the parametric and measurement spaces to demonstrate the manifestation of ion-channel degeneracy in the emergence of signature granule cell physiology. Together, these results reinforce existing lines of evidence for the manifestation of ion-channel degeneracy in DG granule cells, constrained by morphology, by ion-channel gating kinetics and distributions, and several sub- and supra-threshold electrophysiological measurements.

Finally, we used this heterogeneous model population that manifested signature electrophysiological characteristics of DG granule cells to assess forward propagation of synaptic potentials and back-propagation of action potentials. We found the dendritic attenuation characteristics to be comparable with electrophysiological properties of granule cell dendrites, thus validating unconstrained measurements in our model population. Our analyses underscore the critical need to account for ion-channel degeneracy and heterogeneities in granule cells as they play crucial roles in defining their excitability, somato-dendritic and dendro-somatic information transfer, and spatiotemporal summation.

## METHODS

We adapted and retuned a morphologically and biophysically realistic DG granule cell (GC) model from (Beining et al., 2017b) to match 17 different characteristic physiological properties of GCs (Fig. 1) from electrophysiological recordings (Mishra and Narayanan, 2020). The GC morphology was stratified into seven sections: outer molecular layer (OML), middle molecular layer (MML), inner molecular layer (IML), granule cell layer (GCL), soma, axon initial segment (AIS), and axon. The stratification was implemented to non-homogeneously distribute passive and active components across different sections (Table 1), similar to the original model (Beining et al., 2017b). Spines were implicitly accounted for by scaling the leak conductance and specific membrane capacitance in the IML by a factor of 1.45, and in the MML and OML by a factor of 1.9 (Beining et al., 2017b). Leak channels were incorporated across sections non-homogeneously (Hervieu et al., 2001; Gabriel et al., 2002; Yarishkin et al., 2014; Beining et al., 2017b) by altering the leak conductance *g*_p*as*_ (Table 1).

**Figure 1.**
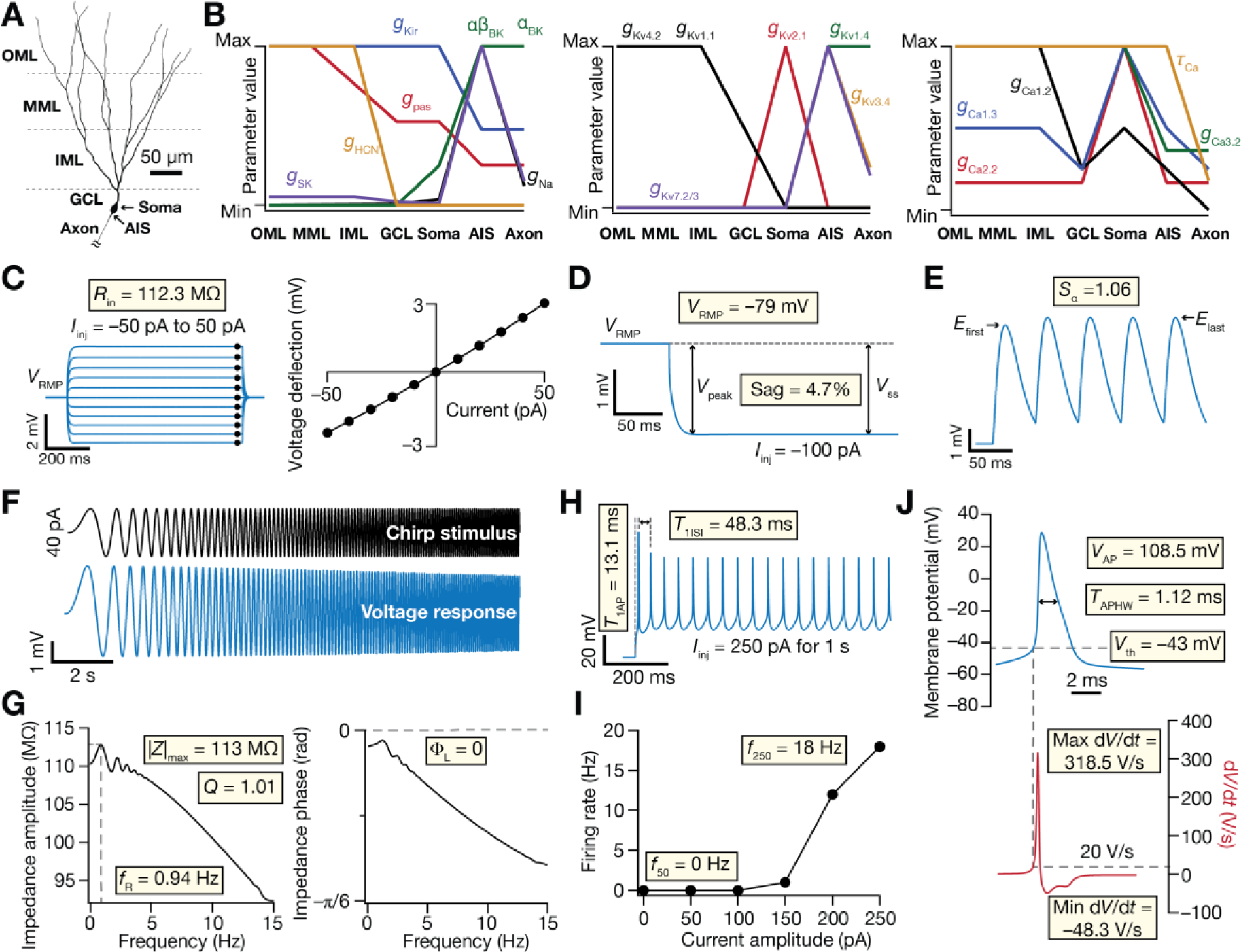
Morphologically realistic granule cell model showing signature physiological measurements. *A*: Two-dimensional projection of the 3D morphology of DG granule cell from (Beining et al., 2017b). OML: outer molecular layer. MML: middle molecular layer. IML: inner molecular layer. GCL: granule cell layer. AIS: axonal initial segment. *B*: Distribution of different ion-channel conductances across the morphological model of granule cells. *C*: *Left*, somatic voltage traces recorded in response to the current injections (*I*_inj_) of – 50 pA to + 50 pA in steps of 10 pA. *Right*, steady-state values of voltage responses from traces in the left plotted against the respective value of injected current. The slope of the linear fit on this plot yielded the input resistance (*R*_in_) of the neuron. *D*: Somatic voltage response to a 100 pA hyperpolarizing current injection. Sag was computed from the steady-state (*V*_SS_) and peak (*V*_peak_) values of voltage deflections from resting membrane potential (*V*_RMP_). *E*: Somatic voltage response to five alpha excitatory postsynaptic current injections to calculate summation ratio (*S*_α_). *F*: *Top*, chirp current stimulus spanning 0– 15 Hz in 15 s of 40 pA peak-to-peak amplitude. *Bottom*, somatic voltage response to chirp current injection. *G*: Impedance amplitude (left) and phase (right) profiles obtained from the chirp current stimulus and the associated voltage response in *F*. *H*: Somatic voltage response showing action potential firing elicited by a 250-pA depolarizing current injection. *I*: Plot of action potential firing rate against injected current amplitude, ranging from 0 to 250 pA in steps of 50 pA. *J*: First action potential from the train of spikes elicited by a 250-pA depolarizing current injection. The first derivative of the voltage response, *dV*/*dt* is plotted below. The voltage value at the timepoint where the derivative crossed 20 V/s was measured as threshold voltage *V*_th_.

**Table 1:**
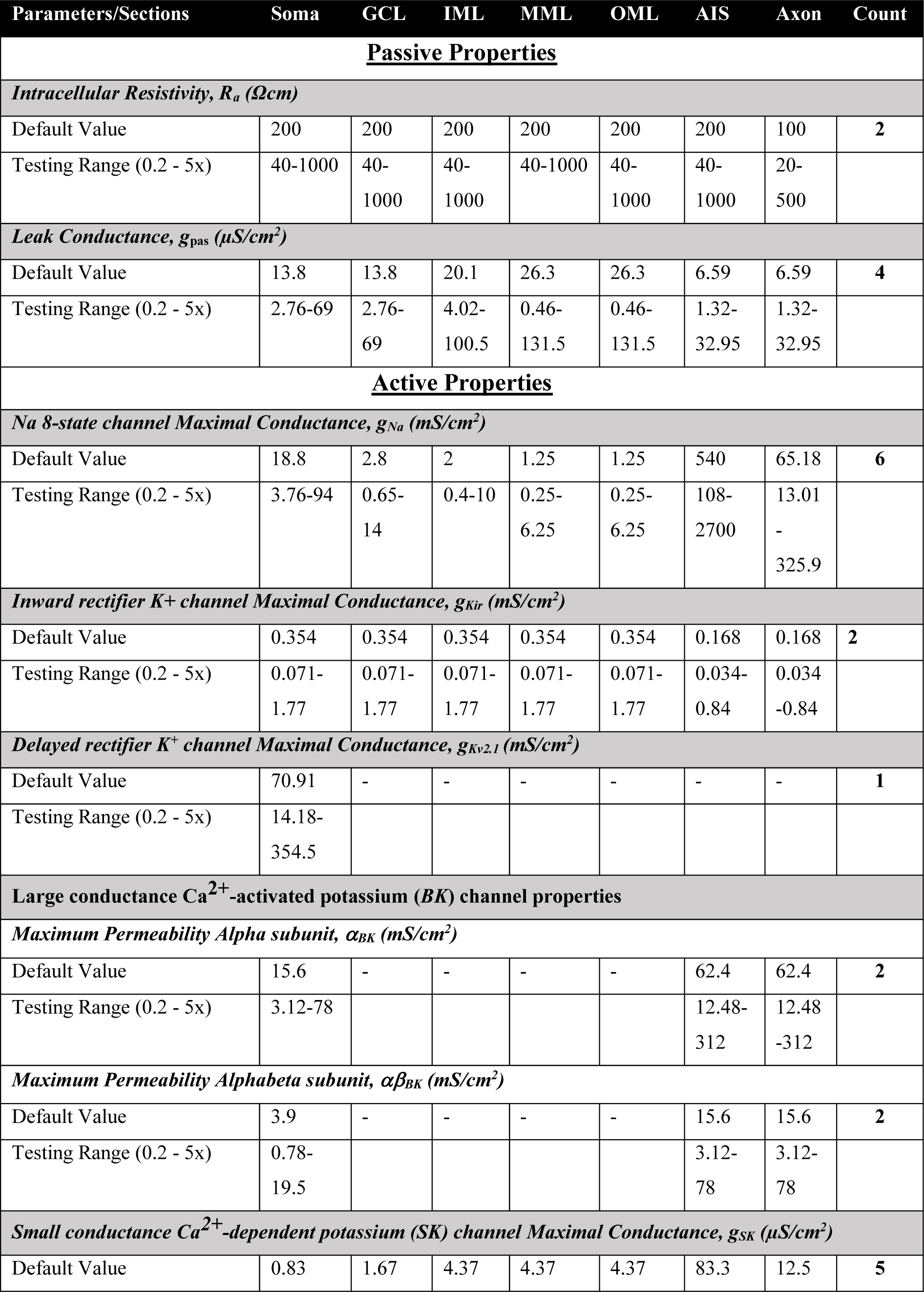
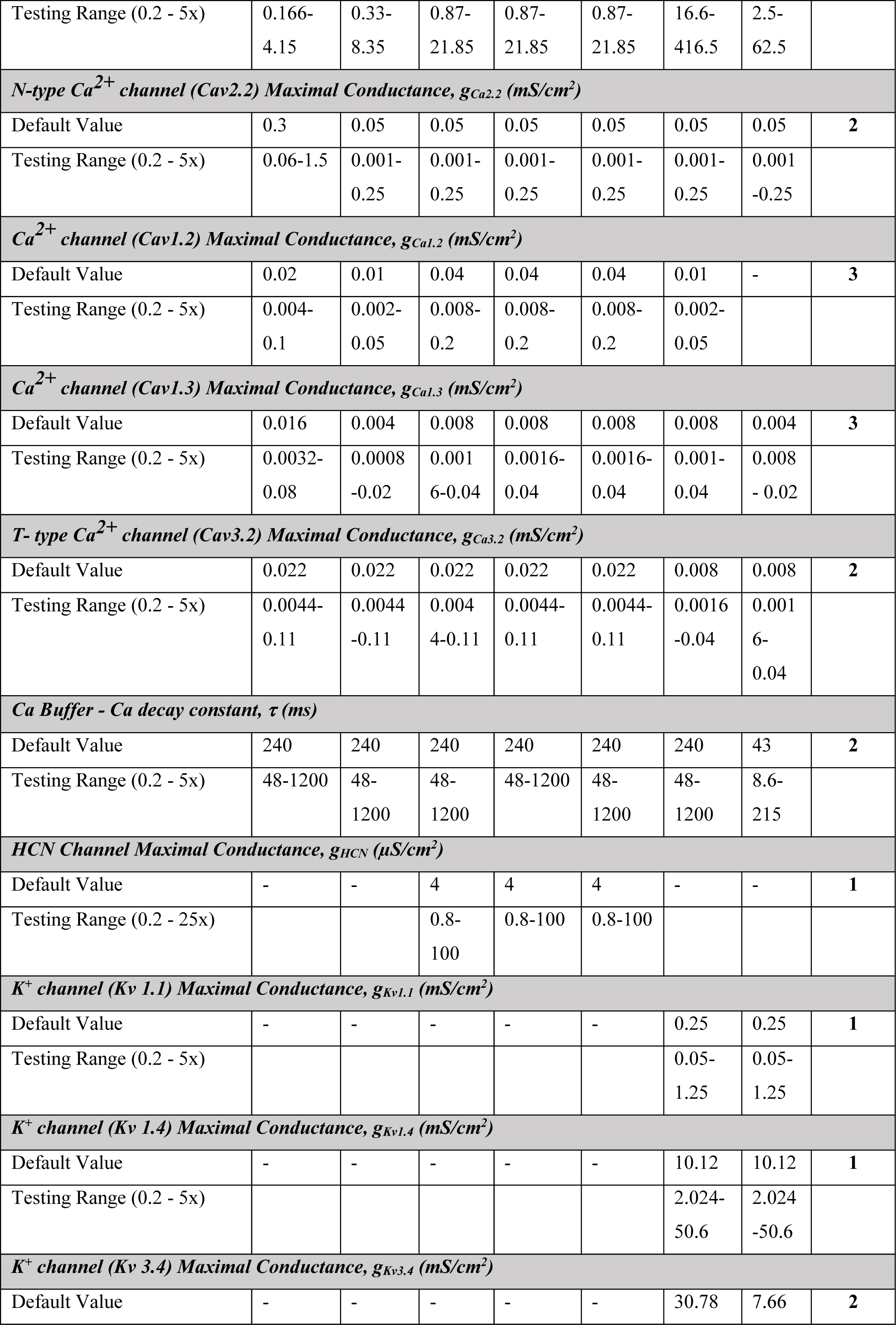
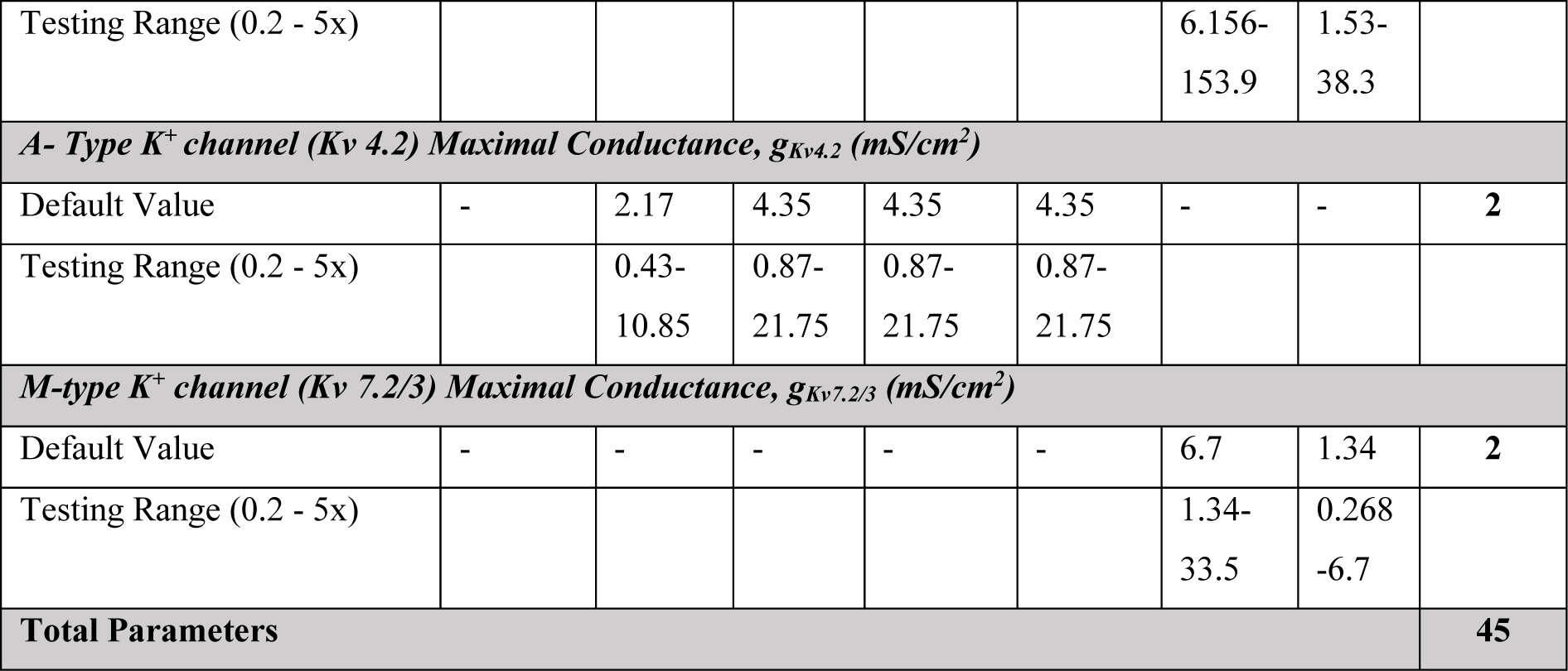
Model parameters with their base values and the range spanned for the stochastic search algorithm for valid DG GCs. All channel names and their gating kinetics were taken from Beining et al. (Beining et al., 2017b)

The 15 active conductances incorporated into the model, with their gating kinetics and distributions adapted and retuned from the original model (Beining et al., 2017b) (Table 1; Fig. 1*B*). An inward-rectifier potassium channel (*K*_ir_) (Lopatin et al., 1995; Yan and Ishihara, 2005; Panama and Lopatin, 2006; Ishihara and Yan, 2007) was set to be present in all sections, with location-dependent distribution of conductance. An 8-state sodium channel (Na) with region-dependent densities was incorporated into the model with the highest density in the AIS and lower densities in dendrites and soma (Kress et al., 2010; Schmidt-Hieber and Bischofberger, 2010). Inactivating voltage-gated potassium channels K_v_1.1 (Christie et al., 1989) and K_v_1.4 (Wissmann et al., 2003) were present in AIS and axon, whereas K_v_4.2 (Barghaan et al., 2008) was localized to dendrites. The delayed-rectifier K_v_3.4 channels (Rudy et al., 1991; Schroter et al., 1991; Rettig et al., 1992; Vega-Saenz de Miera et al., 1992; Riazanski et al., 2001) were incorporated only into the axonal and AIS compartments. *M*-type potassium channels (K_v_7.2/7.3) (Mateos-Aparicio et al., 2014) were localized to the axon and the AIS. The hyperpolarization-activated cyclic nucleotide gated (HCN) non-specific cationic channels (Stegen et al., 2012) were inserted in the dendrites. Voltage-gated *N*-type (Ca_v_2.2) (Fox et al., 1987) and *T*-type calcium channels (Ca_v_3.2) (Burgess et al., 2002) were distributed across all compartments. Voltage-gated *L*-type calcium channels (Ca_v_1.2/1.3) (Evans et al., 2013) were inserted such that Ca_v_1.3 was present in all compartments and Ca_v_1.2 spread across all sections except for the axon. The calcium-dependent big-conductance potassium (BK) channels (Jaffe et al., 2011) were inserted into the soma and axon. The calcium-dependent small-conductance potassium (SK) channels (Hirschberg et al., 1999; Solinas et al., 2007) were incorporated in all the sections.

A majority of these ion channels were modelled using Hodgkin-Huxley dynamics (Hodgkin and Huxley, 1952). *A*-type K_v_4.2 potassium channels followed a 15-state Markovian model. *T*-type Ca_v_3.2 calcium and sodium channels were modelled as 8-state Markovian models. SK and K_ir_ channels were both modelled as 6-state Markov models. Calcium buffer shell model was modified from (Anwar et al., 2014). The Ca^2+^ decay time constant was set to 43 ms (Jackson and Redman, 2003) in the axon and 240 ms (Stocca et al., 2008) in all other compartments. Sodium reversal (*E*_Na_) was set to 50 mV and potassium reversal (*E*_K_) to – 80 mV. Sodium channels (Na8st) were introduced in the dendritic compartments in GCL, IML, MML and OML strata to accommodate active dendrites in GCs (Krueppel et al., 2011). All model parameters and their respective base values are listed section wise in Table 1.

### Sub-threshold measurements

DG GC models were validated against 17 electrophysiological signature characteristics that were measured experimentally (Mishra and Narayanan, 2020). The measurements were computed using well-established procedures (Narayanan and Johnston, 2007; Narayanan and Johnston, 2008; Rathour and Narayanan, 2014; Basak and Narayanan, 2018; Mishra and Narayanan, 2020; Mishra and Narayanan, 2021a) as described below. Resting membrane potential (*V*_*RMP*_) was measured as the potential at which the membrane rest when no current is injected. *V*_*RMP*_ was calculated as the mean of recorded voltage for the last 50 ms of a 1-s simulation performed in the absence of current injection. All sub- and supra-threshold measurements were performed after an initial delay of 1 s to allow *V*_*RMP*_ to reach steady-state value. Input resistance (*R*_*in*_) was measured as the slope of a linear fit to the steady-state *V– I* plot obtained by injecting sub-threshold current pulses of amplitudes spanning – 50 to + 50 pA, in steps of 10 pA (Fig. 1*C*). Percentage sag was measured from the voltage response of the cell to a hyperpolarizing current pulse of – 100 pA for 1000 ms and was defined as: 100[1 – (*V*_*SS*_ / *V*_*peak*_)], where *V*_*SS*_ and *V*_*peak*_ depicted the steady-state and peak voltage deflection from *V*_*RMP*_, respectively (Fig. 1*D*). To assess temporal summation, five α -excitatory postsynaptic currents (α -EPSCs) with 50 ms interval were injected into the somatic compartment. Temporal summation ratio (*S*_*α*_) was computed as *E*_*last*_ /*E*_*firstt*_, where *E*_*last*_ and *E*_*firstt*_ are the amplitudes of last and first α -excitatory postsynaptic potentials, respectively, recorded in response to the injection of 5 α -EPSCs (Fig. 1*E*).

The chirp stimulus used for characterizing the impedance profiles was a sinusoidal current of constant amplitude below firing threshold, with its frequency linearly spanning 0– 15 Hz in 15 s (Fig. 1*F*). The magnitude of the ratio of the Fourier transform of the voltage response (Fig. 1*F*) to the Fourier transform of the Chirp stimulus yielded the impedance amplitude profile (Narayanan and Johnston, 2008):

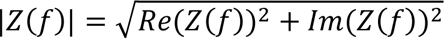

where *Re*(*Z*(*f*)) and *Im*(*Z*(*f*)) were the real and imaginary parts, respectively, of the impedance *Z* as a function of frequency, *f*. The peak value of impedance across all frequencies was measured as the maximum impedance amplitude |*Z*|_m*ax*_. The frequency at which the impedance amplitude reached its maximum value was defined as the resonance frequency (*f*_*R*_). Resonance strength (*Q*) was measured as the ratio of the maximum impedance amplitude to the impedance amplitude at 0.5 Hz. Impedance phase *ϕ*(*f*) was computed as:

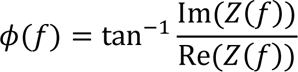

Total inductive phase, Φ_L_, defined as the area under the inductive part of *ϕ*(*f*) (Fig. 1*G*) was defined as (Narayanan and Johnston, 2008):

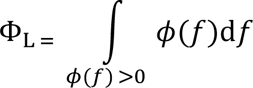

### Supra-threshold measurements

Supra-threshold measurements were obtained through depolarizing current injections, with amplitudes large enough to elicit action potentials (AP), into the cell resting at *V*_*RMP*_ (Mishra and Narayanan, 2019; 2020). AP firing frequency was computed by counting the number of spikes obtained during a 1000 ms current injection (Fig. 1*H*). Current amplitude of these pulse-current injections was varied from 0 pA to 250 pA in steps of 50 pA, to construct the firing frequency *vs*. injected current (*f*– *I*) plot (Fig. 1*I*). Various AP related measurements were derived from the voltage response of the cell to a 250-pA pulse-current injection (Fig. 1*H*; Fig. 1*J*). The temporal distance between the timing of the first spike and the time of current injection was defined as latency to first spike (*T*_1AP_; Fig. 1*H*). The duration between the first and the second spikes was defined as the first inter-spike interval (*T*_1ISI_). AP amplitude (*V*_*AP*_) was computed as the difference between the peak voltage of the first spike and *V*_*RMP*_ (Fig. 1*J*). AP half-width (*T*_APHW_) was the temporal width measured at the half-maximal points of the AP peak with reference to *V*_*RMP*_ (Fig. 1*J*). The maximum (*dV*/*dt*|_*max*_) and minimum (*dV*/*dt*|_*min*_) values were calculated from the temporal derivative of the first action potential obtained with 250-pA current injection (Fig. 1*J*). The voltage in the AP trace corresponding to the time point at which the *dV*/*dt* crossed 20 V/s was defined as AP threshold (*V*_*t*ℎ_) (Fig. 1*J*). The sub- and supra threshold measurements of the base model and their respective experimentally derived bounds are listed in Tables 2 and 3.

**Table 2:** Sub-threshold measurements of base model DG granule cells and their respective electrophysiological bounds derived from Mishra and Narayanan (Mishra and Narayanan, 2020)

|  | Sub-threshold Measurements | Base Model | Lower bound | Upper bound |
| --- | --- | --- | --- | --- |
| 1. | Resting membrane potential, $V_{RMP}$ (mV) | -79.93 | -80 | -70 |
| 2. | Input resistance, $R_{in}$ (MΩ) | 112.34 | 90 | 300 |
| 3. | Maximal impedance amplitude, $\vert Z\vert _{max}$ (MΩ) | 112.81 | 90 | 225 |
| 4. | Resonance frequency, $f_R$ (Hz) | 0.937 | 0.4 | 1.2 |
| 5. | Resonance strength, $Q$ | 1.012 | 1 | 1.2 |
| 6. | Total Inductive Phase ($\Phi_L$) | 0 | 0 | 0.03 |
| 7. | Sag (%) | 4.7 | 1 | 7 |
| 8. | Summation ratio of αEPSPs, $S_\alpha$ | 1.06 | 0.9 | 1.5 |

**Table 3:**
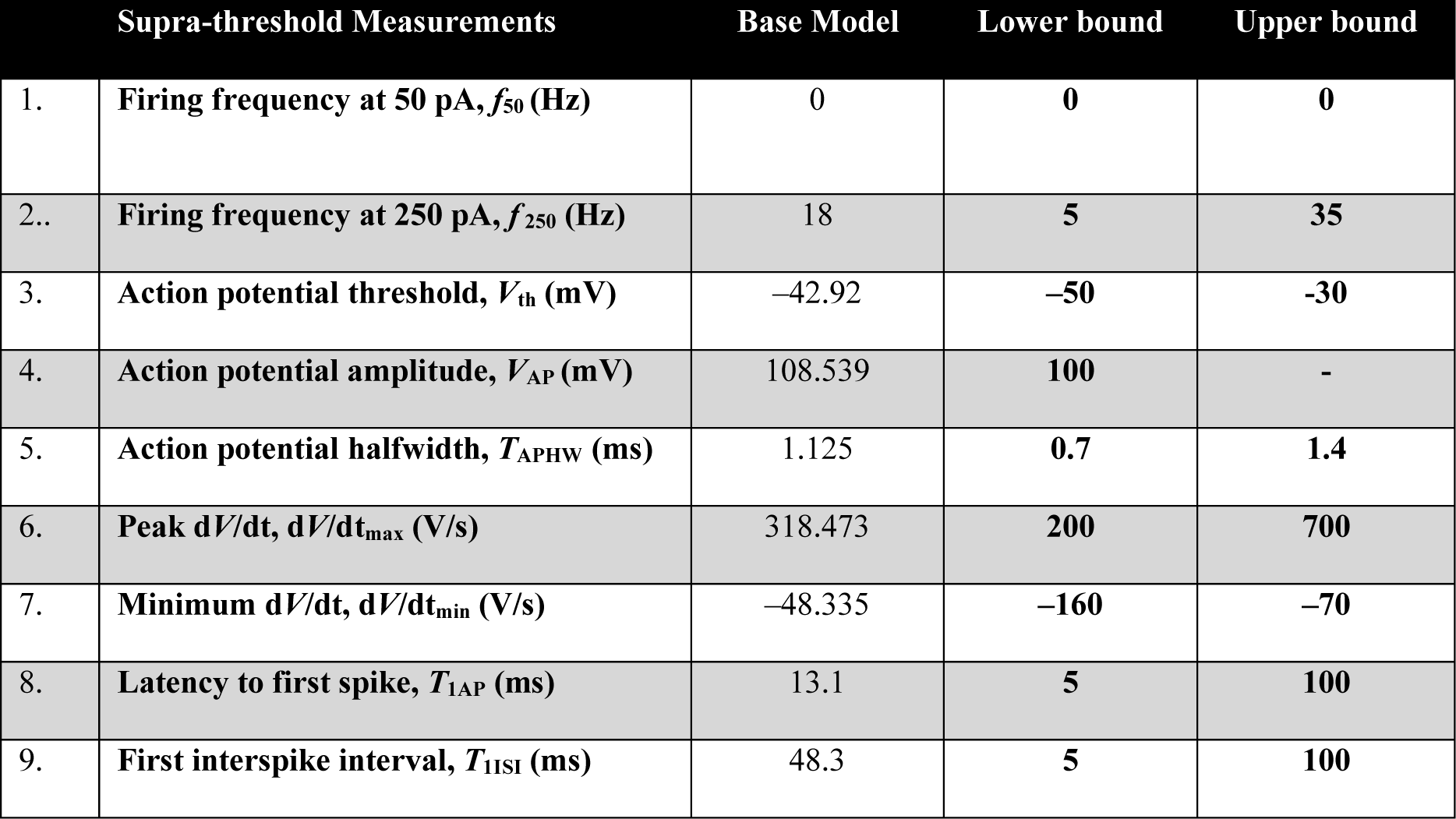
Supra-threshold measurements of base model DG granule cells and their respective electrophysiological bounds derived from Mishra and Narayanan (Mishra and Narayanan, 2020)

### Multi-parametric multi-objective stochastic search (MPMOSS)

We employed multi-parametric multi-objective stochastic search (MPMOSS) to generate a heterogeneous population of GC neuronal models. A randomized search involving a parametric space of 45 dimensions (Table 1) was performed to generate valid models of GCs. Parameters whose distributions were non-identical across sections were split into multiple parameters that depended on the section where they were placed. For example, *K*_ir_ channel has the same conductance value in the soma, GCL, IML, MML, and OML, but a different value for the axon and AIS. Thus, two parameters were defined for *K*_ir_ conductance, ***g***_***kir***_**1** (soma, GCL, IML, MML, OML) and ***g***_***kir***_**2** (axon and AIS).

The base model was hand tuned to match most GC physiological characteristics (Fig. 1; Tables 1– 3). A total of 15,000 random morphologically realistic models were generated by sampling each base model parameters from respective uniform distributions that typically spanned 0.5– 2× of their base values (Table 1). Sub- and supra threshold physiological measurements of each model were computed (Fig. 1) and were validated against their respective bounds obtained from electrophysiological recordings from DG GCs (Tables 2– 3). Models that satisfied all the 17 intrinsic measurement bounds (Tables 2– 3) were declared valid. The parameters and measurements of these valid models were then subjected to further analyses exploring heterogeneities and degeneracy. Pairwise Pearson’ s correlation coefficient (*R*) was calculated across parameters and measurements from the valid models.

### Synapse model and assessment of backpropagating action potentials

Glutamatergic AMPAR synapses were placed in different model compartments with following characteristics of granule cells (Ye et al., 2005; Krueppel et al., 2011). The ionic current through these receptors were modelled using the Goldman– Hodgkin– Katz (GHK) convention (Goldman, 1943; Hodgkin and Katz, 1949; Narayanan and Johnston, 2010). The intra- and extra-cellular concentrations for the different ions were set as: [*Na*]_i_ = 18 mM, [*Na*]_o_ = 140 mM, [*K*]_i_ = 140 mM, and [*K*]_o_ = 5 mM. These ionic concentrations ensured that the reversal potentials for AMPA receptors was set at 0 mV. The AMPAR current was modeled following the GHK convention, and was driven by sodium and potassium:

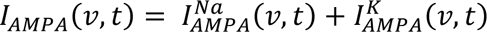

where,

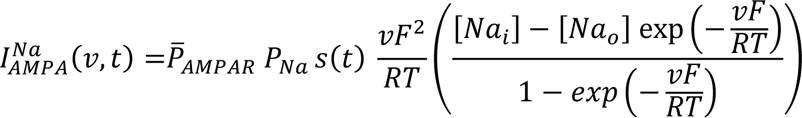

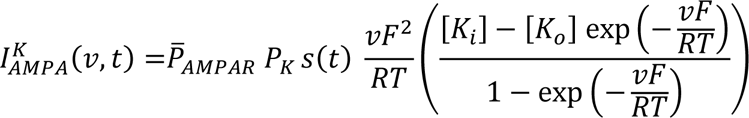

where 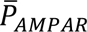 defined the maximum permeability of the AMPA receptors, with *P*_Na_ = *P*_K_ = 1, *R* represented gas constant, and *T* was temperature in Kelvin. *s*(*t*) governed the kinetics of the AMPA receptor current as follows:

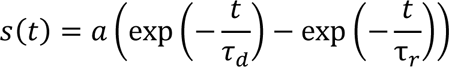

where *a* defined a normalization factor that ensured that 0 ≤ *s*(*t*) ≤ 1. The rise and decay time constants of AMPAR were τ _*r*_ (= 2 ms) and *τ*_*d*_ (= 10 ms) (Ye et al., 2005). The AMPAR density (permeability value 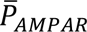) of individual synapses in the base model were adjusted (Narayanan and Chattarji, 2010; Basak and Narayanan, 2018; 2020; Roy and Narayanan, 2021) such that the propagated somatic EPSP amplitude, irrespective of dendritic location, was in the 0.2– 0.3 mV range to match with unitary somatic EPSP amplitudes in DG granule cells (Krueppel et al., 2011). The same location-dependent density values were used across all valid GC models to assess heterogeneities in synaptic information transfer within granule cells.

Backpropagation of action potentials was assessed by initiating a single action potential at the soma and measuring the amplitude at different locations along the dendritic arbor. The recorded amplitudes from all compartments, for each valid GC, were analyzed to assess heterogeneities in backpropagation of action potentials.

### Computational details

All simulations were performed using the NEURON 7.2 programming environment (Hines and Carnevale, 1997) at 34° C. The simulation step size was set as 25 μs. As simulations were associated with morphologically realistic models with a large set of ion channels and associated mechanisms, the computational cost of each simulation for each measurement (especially ones involving large time ranges, such as those for impedance measurements involving a 15 s chirp stimulus on a 25 µs step size) was high. Dimensionality reduction analyses on the measurements and the parametric spaces were performed with principal component analysis (PCA), *t*-distributed stochastic neighbor embedding, *t*-SNE (Van der Maaten and Hinton, 2008); uniform manifold approximation and projection, UMAP (McInnes et al., 2018); and potential of heat-diffusion for affinity-based trajectory embedding, PHATE (Moon et al., 2019). Data analysis and plotting of graphs were done using custom-built software written in MATLAB or Igor Pro programming environment (WaveMetrics Inc., USA).

## RESULTS

The aim of this study was to assess ion-channel degeneracy in morphologically realistic models of DG granule cells, which were constrained by a comprehensive set of sub- and supra-threshold electrophysiological measurements acquired in the laboratory (Mishra and Narayanan, 2020). Towards this goal, we first adapted a detailed morphological model of granule cells from (Beining et al., 2017b), whose biophysical properties (ion channel gating kinetics, distributions, and calcium handling mechanisms; Table 1) were derived from DG granule cells. We retuned this model to account (Fig. 1) for 17 different sub- and supra-threshold electrophysiological measurements recorded from granule cells (Mishra and Narayanan, 2020). We introduced several changes to channel conductances and other properties in adapting the original model (Table 1). An important difference in our model was to account for active propagation of action potentials within granule cell dendrites (Krueppel et al., 2011) by incorporating spike-generating conductances into dendritic compartments. The hand-tuned base model matched the ranges of most electrophysiological measurements (Fig. 1) and formed a substrate for generating a population of DG granule cells.

### A small subset of randomly generated models satisfied all signature electrophysiological characteristics of granule cells

To avoid biases associated with using a single hand-tuned model, we employed a multi-parametric multi-objective stochastic search (MPMOSS) algorithm over a large parametric space to identify valid GC models. The 45-dimensional parametric space spanned all ion-channel conductances and calcium handling across all spatial locations within the morphologically realistic model (Table 1). A total of 15,000 models were generated randomly by sampling independent uniform distributions associated with the 45 parameters, each spanning respective bounds (Table 1). Among these randomly generated models, valid GC models were those that satisfied all 17 sub-(Table 2) and supra-threshold (Table 3) electrophysiological measurements from granule cells (Mishra and Narayanan, 2020). Each measurement required a different protocol and stimulus (or stimuli), which matched with their respective electrophysiological counterparts (Mishra and Narayanan, 2020). Of the 15,000 random models, we found 141 (0.94%) to satisfy all 17 validation criteria (Tables 2– 3). Thus, while arbitrary random combinations of the 45 parameters did not yield valid GC models (> 99% models were invalid), there was a small subset of such combinations that yielded valid GC models. In what follows, we analyze the measurements and parameters associated with this subset of valid models to assess various aspects of their biophysical and physiological characteristics.

### Pronounced heterogeneities in and weak cross-dependencies between measurements from valid granule cell models matched with their electrophysiological counterparts

All 17 electrophysiological measurements were plotted for the 141 valid models (Fig. 2*A*) to assess if they are clustered or were distributed across the range of their respective validation bounds (Tables 2–3). We found the 141 valid GC models to manifest heterogeneous physiological measurements (Fig 2*A*), reflecting the heterogeneities observed in their biological counterparts (Mishra and Narayanan, 2020). Expectedly, the firing rate for a 50-pA current injection, *f*_50_, was identically zero for all models and two impedance measurements (resonance frequency, *f*_*R*_ and total inductive phase, Φ_*L*_) were clustered with low values (Mishra and Narayanan, 2020). The low-pass nature of the granule cell impedance profile translates to low resonance frequency values and minimal inductive phase (Mishra and Narayanan, 2020), thus resulting in clustered values for these measurements (Fig 2*A*).

**Figure 2.**
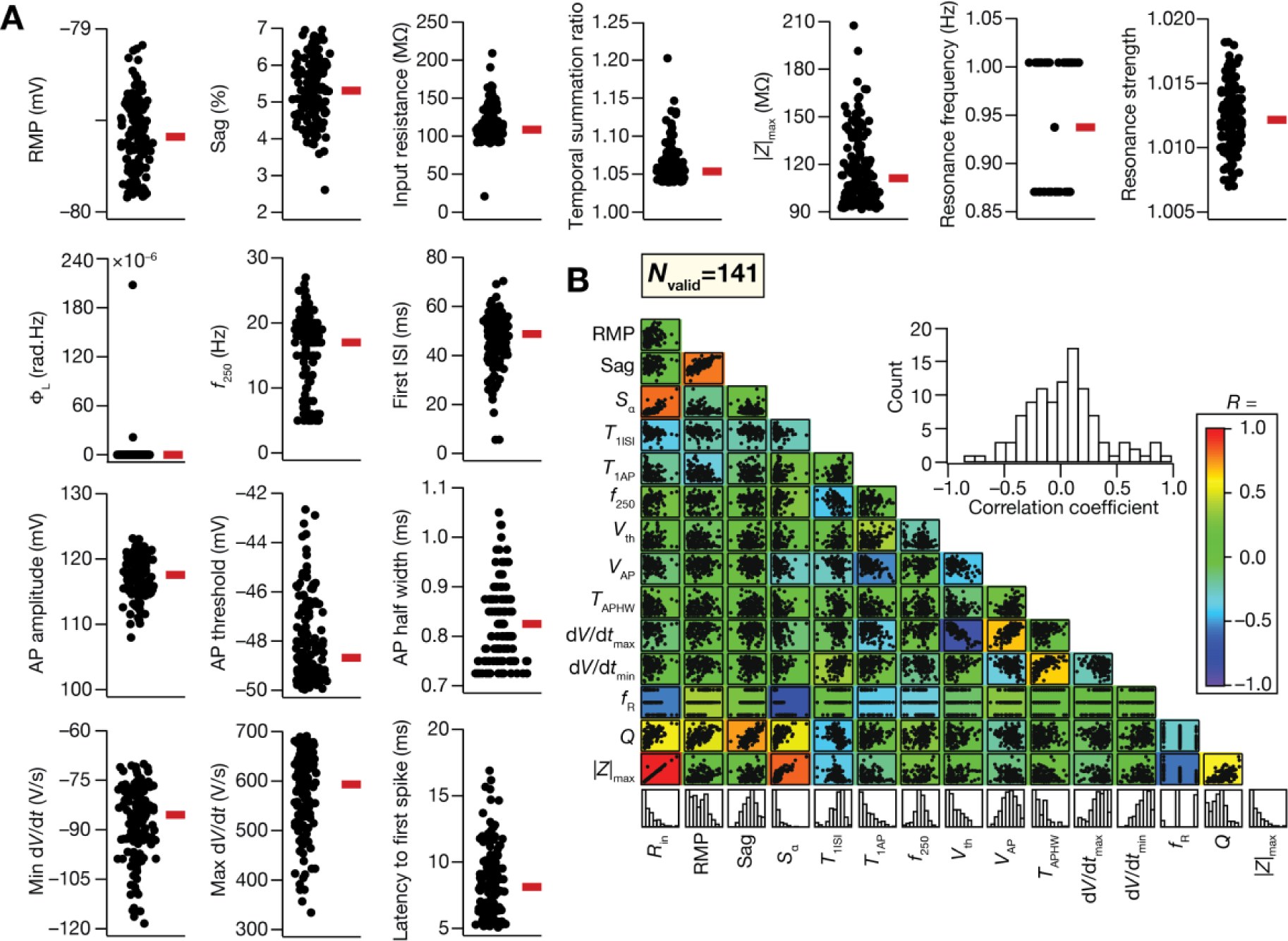
Heterogeneous distribution of characteristic physiological measurements in the valid granule cell models obtained by MPMOSS. (A) Bee-swarm plots depicting the distribution of 8 sub-threshold and 8 supra-threshold measurements of the 141 valid models. The red rectangle adjacent to each plot depicts the respective median value. Of the 17 measurements used for validation, *f*_50_ for all valid models was identically zero by virtue of validation requirements (Table 2) and therefore is not shown. (B) Lower triangular matrix representing pair-wise correlations between 15 measurements underlying all valid models. Each box in the matrix depicts the scatter plot between the respective pair of measurements. The bottommost row represents the histograms of individual measurements in the valid model population. A heat map of Pearson’ s correlation coefficient values corresponding to each scatter plot is superimposed on the matrix. *Inset*, Histogram of the correlation coefficients spanning all pairs. Firing frequency at 50 pA (*f*_50_) and total inductive phase (Φ_L_) were not considered for the pairwise analysis as they were either identically zero for all models (*f*_50_) or had very low values in a few models with the rest measured as zero (Φ_L_).

Turning to cross-dependencies across measurements in valid models, we asked if there were strong pair-wise correlations between these measurements. Strong pairwise correlations would either correspond to similar dependencies on the different ion channels or imply that the different measurements were not qualitatively distinct from each other and were capturing the same physiological characteristics. While multiple measurements might be used to constrain models, a wide set of strong pairwise correlations between these measurements would translate to insufficient constraints on the model validation process. We found a large majority of the pairwise Pearson’ s correlation values to be weak (between – 0.4 to 0.4, defined as weak as per existing descriptions (Evans, 1996)) and non-significant (Fig. 2*B*). A small percentage of measurement pairs showed strong correlations, in a manner that was consistent with electrophysiological measurements from granule cells (Mishra and Narayanan, 2020). Specifically, we found strong positive correlations between *R*_in_, |*Z*|_max_, and *S*_α_, similar to their dependencies in measurements from rat granule cells (Mishra and Narayanan, 2020). This was expected because these sub-threshold measurements of excitability are dependent on same sets of passive and active properties. We found strong negative correlation between *dV*/*dt*|_max_ and *V*_th_, which was expected because the voltage threshold will be more hyperpolarized if the rate at which voltage rises towards 20 V/s were high (used in computing *V*_th_; Fig. 1*J*).

We applied linear and nonlinear dimensionality reduction techniques on the measurement space associated with the 141 valid models, the outcomes of which also did not suggest strong cross-dependencies across the different measurements in valid models (Fig. 3). Together, the lack of strong dependencies across measurements (Fig. 2*B*, Fig. 3) demonstrate that these measurements were quantifying disparate aspects of granule cell physiology. In summary, our model population to be representative of the rat granule cell population because of the consistent relationship between our model population and experimental findings (Mishra and Narayanan, 2020) with reference to: (a) individual measurements (Fig. 2*A*); (b) their pairwise co-dependencies (Fig. 2*B*); and (c) the lack of strong structure in the reduced dimensional measurement space (Fig. 3).

**Figure 3.**
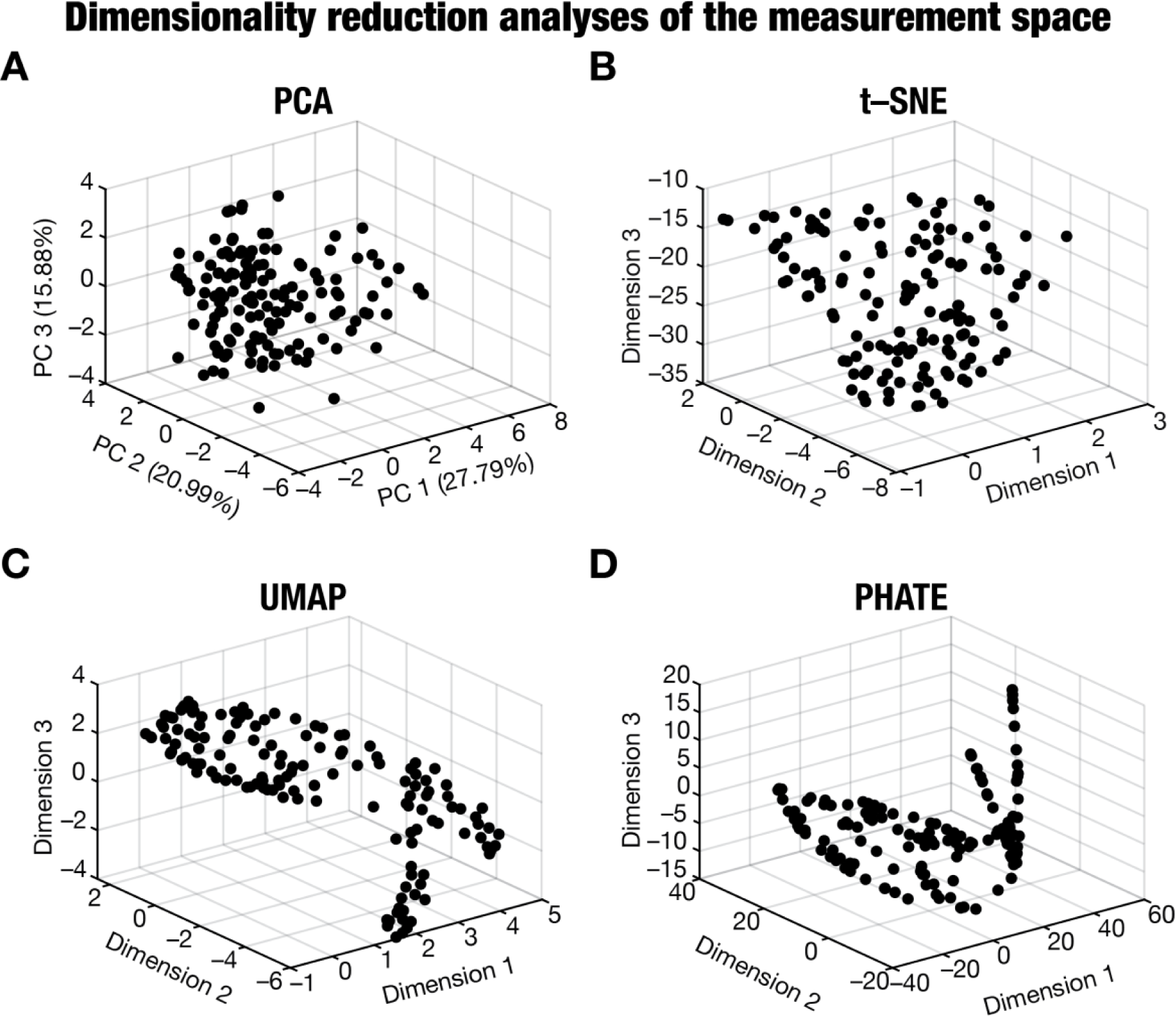
Linear and non-linear dimensionality reduction analyses on the measurement space of the valid granule cell models. *A– D*: Outcomes of principal component analysis, PCA (A), *t*-distributed stochastic neighbor embedding, *t*-SNE (B), uniform manifold approximation and projection, UMAP (C), and potential of heat-diffusion for affinity-based trajectory embedding, PHATE (D) on the 15-dimensional measurement space of the 141 valid granule cell models. The axes labels in panel A depict the percentage variance explained by the respective principal component (PC). Firing frequency at 50 pA (*f*_50_) and total inductive phase (Φ_L_) were not considered for the pairwise analysis as they were either identically zero for all models (*f*_50_) or had very low values in a few models with the rest measured as zero (Φ_L_), thus yielding a 15-dimensional space of the 17 measurements considered in Table 1.

### Ion-channel degeneracy and weak parametric cross-dependencies in valid granule cell models

A small subset of random parametric combinations yielded valid granule cells models that satisfied all characteristic physiological properties. We analyzed the specific distributions of parameters and their cross-dependencies in these valid models. We noted that > 99% random models did not satisfy all physiological constraints. This rules out one extreme of the randomness continuum, where any arbitrary set of random values assigned to these parameters would not yield valid models. The other extreme is a scenario where there is a completely determined, single parametric combination that satisfies all physiological constraints, and all 141 valid models involve small shifts around that single parametric combination. To assess this scenario where all valid models are clustered around a single valid parametric combination, we first picked 6 of the 141 valid models that had very similar sub-(Fig. 4*A– B*) and supra-threshold (Fig. 4*C*) physiological measurements. We plotted the 45 parameters associated with these functionally similar models spanning their respective min-max ranges from Table 1 (Fig. 4*D*). We found that the parameters associated with these six valid models spanned a large range of their min-max ranges (Fig. 4*D*). We looked at the distribution of each parameter across all 141 valid models that manifested characteristic GC physiology (Fig. 5). We found all of them to cover a wide span of their respective min-max ranges. Thus, there were several specific ion-channel combinations, with each parameter spanning a wide range, that can satisfy all constraints associated with granule cell physiology. Together, these analyses rule out the other extreme of the randomness continuum, whereby all 141 models thus were clustered around a completely determined single parametric combination.

**Figure 4.**
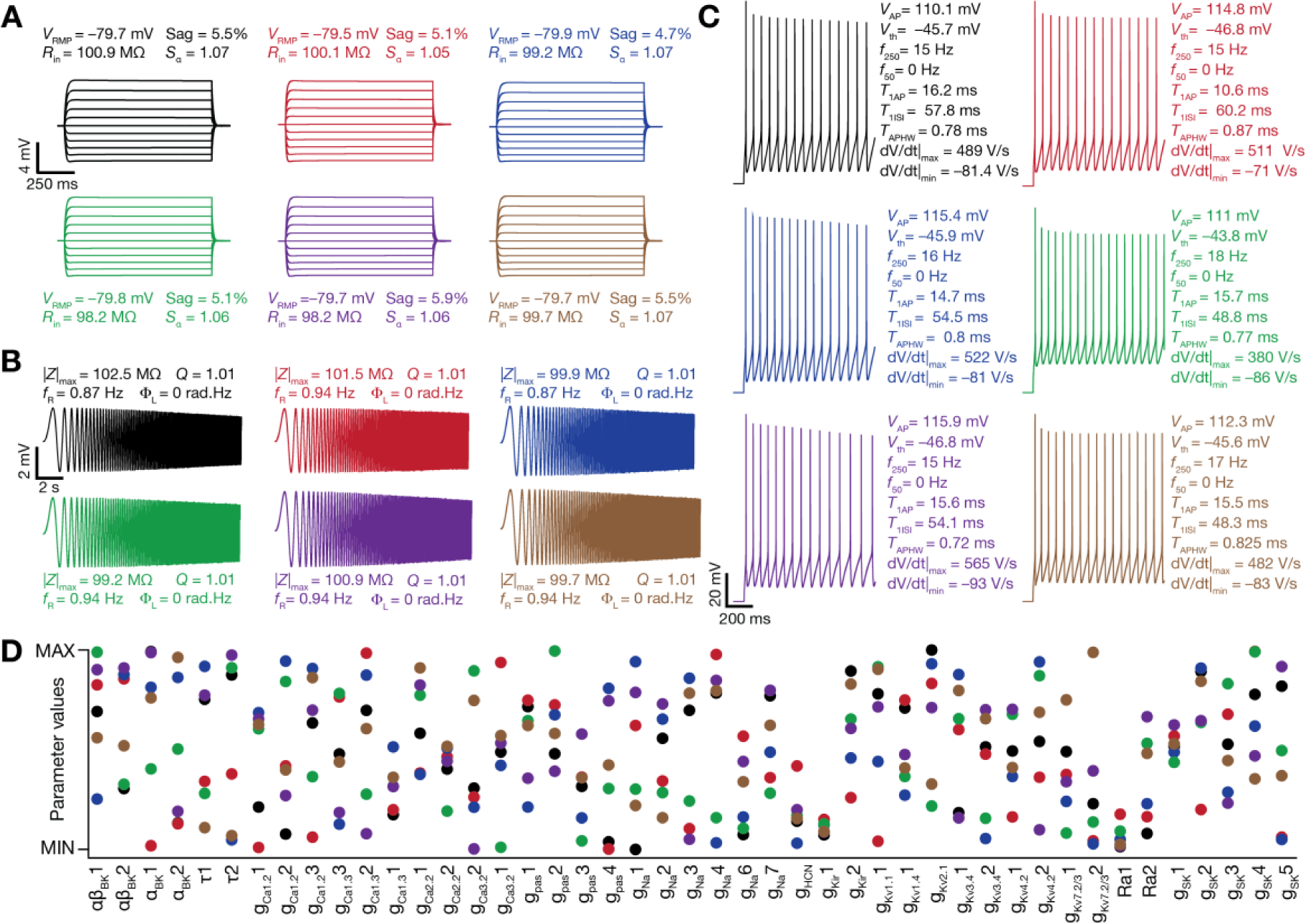
Disparate combinations of model parameters yielded similar physiological measurements in 6 randomly chosen valid granule cell models. *A*: Voltage responses of 6 different valid models with similar measurements for current injections spanning – 50 to + 50 pA in steps of 10 pA. *B*: Voltage responses of the six functionally similar models to chirp current injection. *C*: Voltage responses of the six models to a 250-pA depolarizing current injection showing action potential firing. All 17 measurements for the 6 similar models are depicted across panels *A–C*. *D*: Normalized values of the 45 parameters that defined each valid GC model, shown for the 6 functionally similar models whose measurements are depicted in panels *A–C*. Normalization was with reference to the respective minimum and maximum bounds for that parameter (Table 1). Distinct colors uniquely identify different models across all panels.

**Figure 5.**
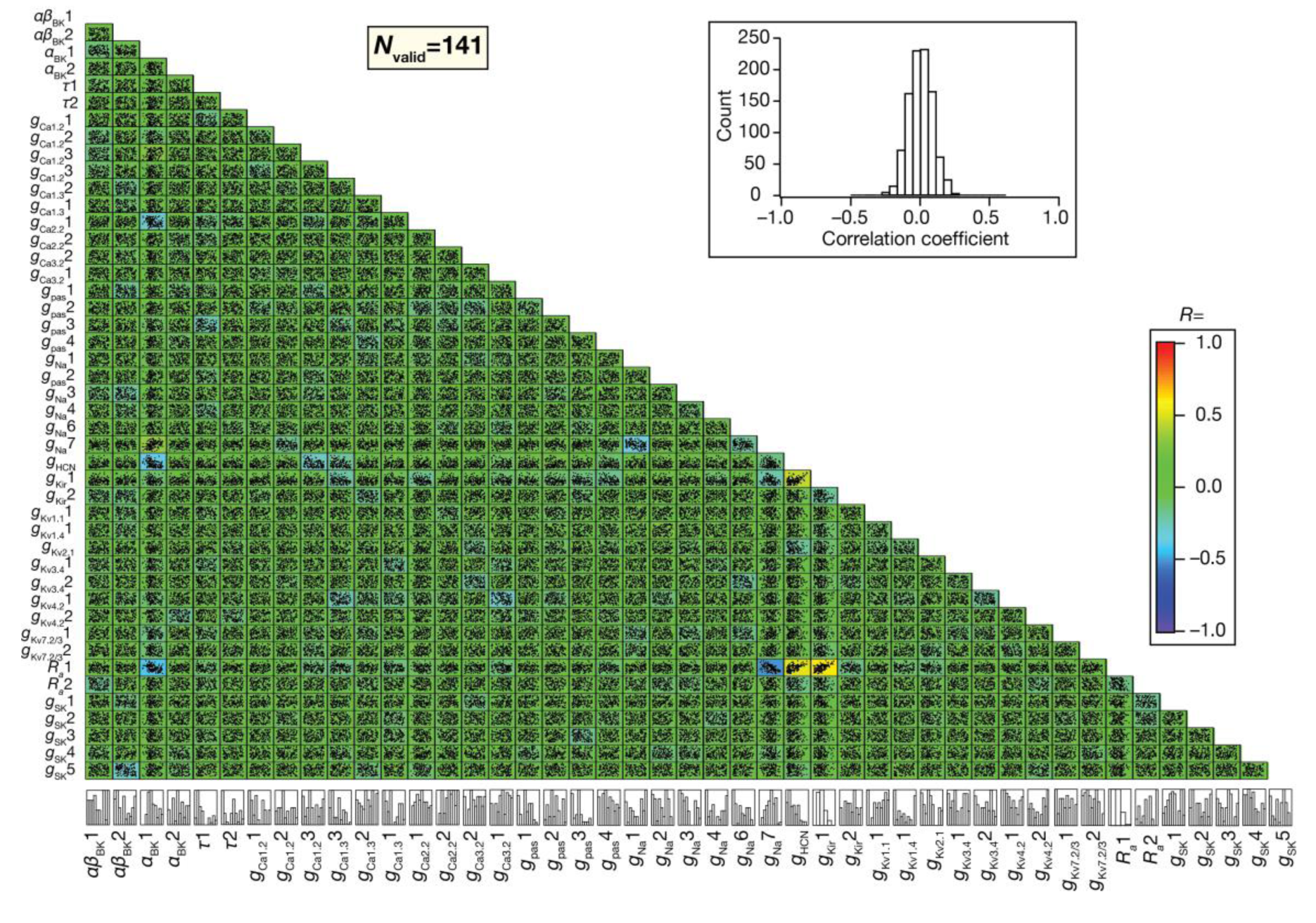
Weak pairwise correlations between parameters that defined the valid granule cell model population. Lower triangular matrix representing correlation between the values of the 45 parameters that defined the 141 valid models. Each box in the matrix depicts the scatter plot between respective parameter pairs. The bottommost row represents the histograms for individual parameters in the valid model population. A heat map of Pearson’ s correlation coefficient values corresponding to each scatter plot is superimposed on the matrix. *Inset*, Histogram of the correlation coefficients spanning all pairs.

We assessed if the absence of clustering in the parametric space was because of strong parametric dependencies, where reduction in one parametric value was compensated in a pairwise manner by either increase or decrease in the value of another parameter. Such pairwise relationships could occur if two different ion channels were functionally similar where the one could be replaced by another without loss of function. In addition, there could be functions that require pairwise increase or decrease of specific pairs of ion channels, which could also be contributing such pairwise relationships. A widespread prevalence of such pairwise relationships would imply that the parametric space was not as high-dimensional as the numbers indicate but reduces to a low-dimensional space where highly cross-dependent parameters covary to yield valid models. To assess such pairwise relationships, we first computed Pearson’ s correlation coefficients between each unique pair of parameters in the valid models. We found that none of the parametric pairwise correlations crossed an absolute value of 0.4, indicating very weak to weak correlations (Evans, 1996) between parameters of the valid models (Fig. 5). Thus, there were no strong pairwise relationships between parameters, indicating the absence of strong pairwise cross-dependence in the parametric space.

The absence of pairwise correlations ruled out strong pairwise dependencies. However, strong cross-dependencies involving several parameters could still yield an effectively low-dimensional space associated with valid model parameters. We asked if the parametric space associated with valid models was indeed high-dimensional or was mapped onto a low-dimensional space by applying linear (Fig. 6*A*) and nonlinear (Figs. 6*B– D*) dimensionality reduction techniques on the parametric space. We found the variance explained by each of the first three principal components with PCA (Fig. 6*A*) to be minimal and a lack of a low-dimensional structure with any of the four dimensionality reduction techniques that we employed (Fig. 6*A–D*). Together, the absence of strong pairwise correlations (Fig. 5) and the lack of structured low-dimensional spaces (Fig. 6) ensure the high-dimensional nature of the parametric space and rule out strong cross-dependencies across different model parameters.

**Figure 6.**
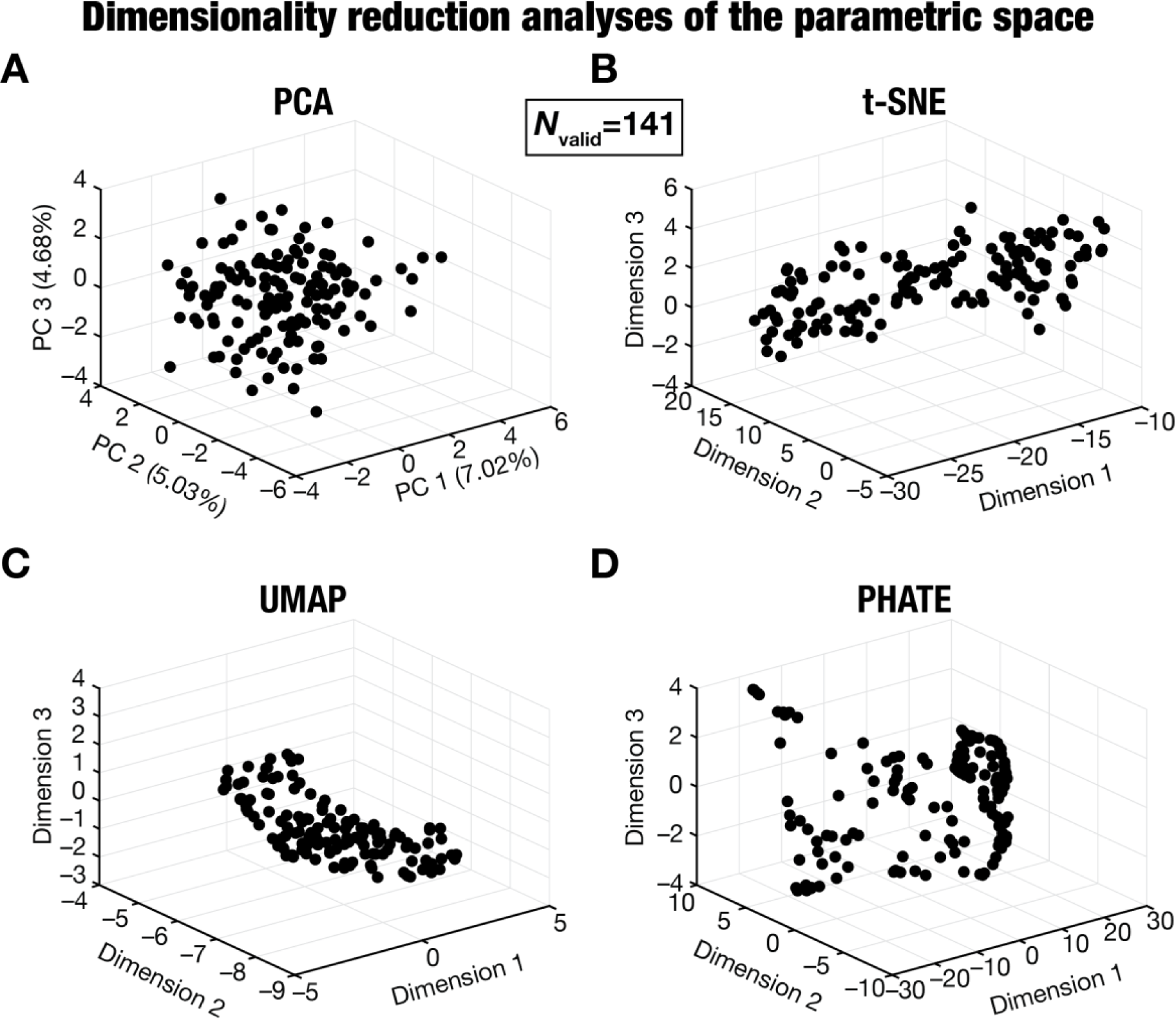
Linear and non-linear dimensionality reduction analyses on the parameter space of the valid granule cell models. *A– D*: Outcomes of principal component analysis, PCA (A), *t*-distributed stochastic neighbor embedding, *t*-SNE (B), uniform manifold approximation and projection, UMAP (C), and potential of heat-diffusion for affinity-based trajectory embedding, PHATE (D) on the 45-dimensional parametric space (Table 2) of the 141 valid granule cell models. The axes labels in panel A depict the percentage variance explained by the respective principal component (PC).

### Heterogeneities in propagation physiology of valid granule cell models

Did all valid granule cell models manifest similar information transfer characteristics across the somato-dendritic arbor? We analyzed synaptic information transfer by studying unitary activation of synapses located at each of the several dendritic locations in the GC morphology (Fig. 7*A*). In the base model, we activated a single synapse containing AMPA receptors at a selected dendritic location, adjusting the receptor density such that the propagated somatic EPSP was in the 0.2– 0.3 mV range to match with electrophysiological recordings from granule cells (Krueppel et al., 2011). We measured the local dendritic voltage at the location of the synapse and the propagated somatic voltage for this location (Fig. 7*B*). We repeated this for all dendritic locations (Fig. 7*C*). As expected, we found that with increasing path distance of the synaptic location from the soma, there was an increase in the local dendritic voltage required to maintain the somatic EPSP within the 0.2– 0.3 mV range (Fig. 7*C*). At the end of this procedure in the base model, we had a value for the AMPAR density for each location across the dendritic arborization. We used these location-dependent receptor density values (obtained from the base model) to assess heterogeneities of propagation across the 141 valid GC models. Specifically, if all valid models behaved the same as the base model, then the somatic EPSP amplitude will be in the 0.2– 0.3 mV for all valid models, irrespective of synaptic location. A deviation from this range would indicate heterogeneities in synaptic information transfer across valid models.

**Figure 7.**
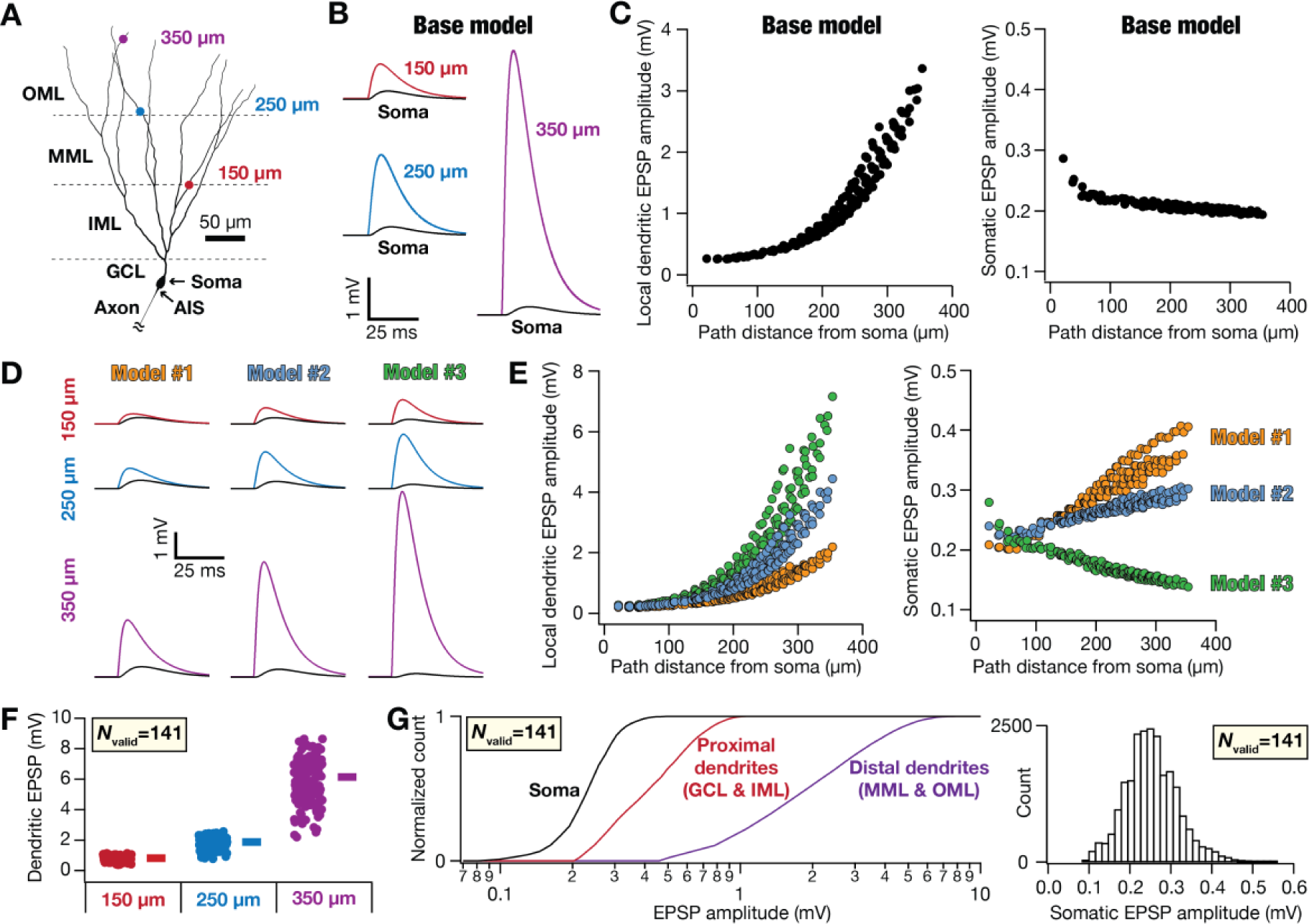
Heterogeneities in forward synaptic propagation across the somato-dendritic arbor of valid granule cell models. *A*: 2D projection of the 3D morphology of DG granule cell showing the 4 locations (Soma, 150 µm, 250 µm, and 350 µm) where illustrative examples of heterogeneities are shown. The numbers represent path distances of marked dendritic locations from the soma. *B*: Excitatory postsynaptic potentials (EPSP) generated by the activation of single synapses located at each of the 3 dendritic locations shown in panel A. The local dendritic voltage trace at the specified synaptic location (colored traces) and the propagated somatic voltage trace (black) are plotted for each of the three dendritic locations on the base model are shown. *C*: Local dendritic EPSP amplitude (*left*) and the propagated somatic counterpart (*right*) plotted against the location of the single synapse on the base model. It may be noted that the somatic voltage was in the 0.2– 0.3 mV range irrespective of synaptic location (*right*). The local dendritic voltage required to retain somatic voltage at 0.2– 0.3 mV increased with increase in distance from the soma (*left*). *D*: Local dendritic EPSP traces and the propagated somatic counterpart (black) for the three synaptic locations (from panel *A*), shown for three different valid models. *E*: Local dendritic EPSP amplitude (*left*) and the propagated somatic counterpart (*right*) plotted against the location of the single synapse for the 3 valid models illustrated in panel D. The heterogeneities in the recorded amplitudes might be noticed across the three models, as well as with reference to the base model (panel *C*) *F*: Local dendritic EPSP amplitude recorded at the 3 dendritic locations marked in panel A for all 141 valid models. *G*: *Left*, Cumulative histograms of somatic and local dendritic EPSP amplitude recorded for all locations for all 141 valid models. Histograms are shown for somatic, proximal dendritic (encompassing GCL and IML), and distal dendritic (locations in MML and OML) locations. *Right*, Histogram of somatic EPSP amplitude for all locations for all 141 valid models. For *D– F*, the AMPAR density of synapses across all valid models were taken from the base model for each synaptic location (panel *C*).

We found pronounced heterogeneities in synaptic information transfer across different valid models. Three illustrative examples are shown in Figure 7*D–E*, demonstrating wide variability in local dendritic synaptic responses as well as in propagated somatic voltages, or an identical synapse at the same location across models. Although the dendritic voltage responses were larger with increased synaptic distance from the soma (Fig. 7*D*), there were differences across models in the magnitude of this increase as well in how they propagated to the soma. These are consequent to differences in active dendritic components of the different models. Importantly, when we assessed local dendritic (Fig. 7*F–G*) and somatic (Fig. 7*G*) voltages across all 141 valid models, we found their ranges to be consistent with their experimental counterparts (Krueppel et al., 2011). Specifically, the somatic unitary EPSP was predominantly in the 0.1– 0.4 range, with a dominant proportion falling within the 0.2– 0.3 mV. The local dendritic voltage was in the 0.2– 0.9 mV for proximal dendritic locations and in the 0.5– 9 mV range for distal dendritic locations (Fig. 7*G*), which match with the ranges reported from electrophysiological recordings (Krueppel et al., 2011). Although synaptic information transfer was not used as a validation criterion for model generation, we found the dendritic unitary synaptic responses and the propagated somatic voltages to match with their respective distributions across rat granule cells.

We assessed backpropagation of action potentials by initiating a single action potential at the cell body and recording the action potential at all somato-dendritic locations. We repeated this for all 141 valid models to assess heterogeneities in action potential backpropagation. We found pronounced heterogeneities in the profile of backpropagation. Whereas some models sustained large-amplitude back-propagating action potentials, others manifested considerable attenuation (Fig. 8*A– D*). The overall reduction in the amplitude of dendritic action potentials (Fig. 8*D– E*) and heterogeneities therein (Fig. 8) were comparable with electrophysiological ranges reported for DG granule cells (Krueppel et al., 2011). Thus, although backpropagating action potentials were not explicitly constrained in the validation process of these models, we found these models to match electrophysiological counterparts in action potential backpropagation and associated heterogeneities across models.

**Figure 8.**
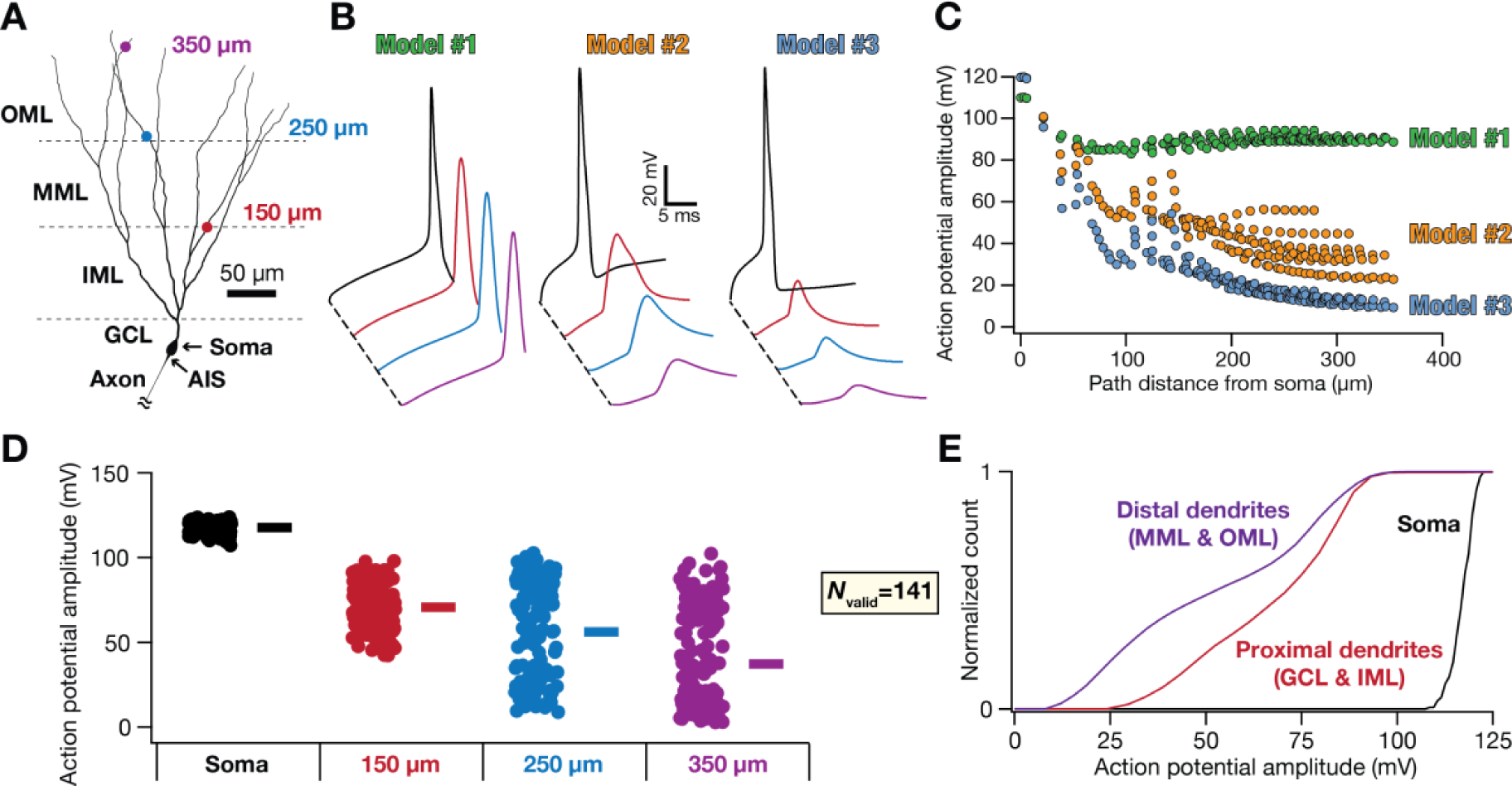
Heterogeneities in backpropagation of action potentials across the somato-dendritic arbor of valid granule cell models. *A*: Two-dimensional projection of the 3D morphology of DG granule cell showing the 4 locations (Soma, 150 µ m, 250 µ m, 350 µ m) where illustrative examples of heterogeneities are shown. The numbers represent path distances of marked dendritic locations from the soma. *B*: Propagating action potential traces of 3 different valid models at the 4 locations marked in panel A. *C*: Action potential amplitude recorded at all somato-dendritic locations for the 3 valid models illustrated in panel B. *D*: Action potential amplitude recorded at the 4 locations marked in panel A for all 141 valid models. *E*: Cumulative histograms of action potential amplitude recorded at all locations for all 141 valid models. Histograms are shown for somatic, proximal dendritic (encompassing GCL and IML), and distal dendritic (locations in MML and OML) locations.

## DISCUSSION

Our study involved three-dimensional, morphologically realistic models of the dentate gyrus granule cell that were constrained by the set of ion channels that they express and several electrophysiological measurements that characterize their cellular neurophysiology. We employed a computationally complex stochastic search algorithm spanning all biophysical parameters that defined the model to search for valid granule cells models. A small subset of valid model, among thousands of randomly generated models, satisfied several sub- and supra-threshold electrophysiological measurements that were obtained from dentate gyrus granule cells (Mishra and Narayanan, 2020). Somato-dendritic measurements from this valid model population were heterogeneous and matched with their electrophysiological counterparts. The different electrophysiological measurements used to identify valid models quantified different aspects of granule cells physiology and matched pairwise dependencies of their electrophysiological counterparts (Mishra and Narayanan, 2020). Importantly, the valid model parameters were neither arbitrarily random nor clustered around a single parametric combination and showed weak cross-dependencies in the parametric space. These observations are consistent with a system that manifests a high degree of complexity, involving specific combinations of disparate ion-channel contributions towards achieving characteristic granule cell physiology. In addition, consistent with complex systems, the emergence of granule cell characteristic physiology showed degeneracy (Edelman and Gally, 2001). Specifically, disparate combinations of functionally segregated subsystems (ion channels and other biophysical systems) yielded similar functional outcomes in the integrated complex system (the granule cell), resulting in the coexistence of functional segregation and functional integration within the complex system (Tononi et al., 1994; Sporns et al., 2000; Edelman and Gally, 2001).

### Heterogeneities and degeneracy in neuronal physiology

Biological complex systems manifest pronounced heterogeneities and exhibit degeneracy in the realization of precise functional outcomes. The nervous system in general, and the mammalian hippocampal formation with specific reference to this study, is no exception to this observation (Tononi et al., 1994; Sporns et al., 2000; Goldman et al., 2001; Price and Friston, 2002; Prinz et al., 2004; Leonardo, 2005; Gjorgjieva et al., 2016; Rathour and Narayanan, 2019; Goaillard and Marder, 2021; Mishra and Narayanan, 2021d; Seenivasan and Narayanan, 2022; Marom and Marder, 2023). Within the mammalian hippocampal formation, degeneracy in the emergence of characteristic physiological properties has been observed in pyramidal neurons of the CA1 (Rathour and Narayanan, 2012; 2014; Srikanth and Narayanan, 2015; Rathour et al., 2016; Migliore et al., 2018; Jain and Narayanan, 2020; Srikanth and Narayanan, 2023), pyramidal neurons of the CA3 (Roy and Narayanan, 2023), basket (Mishra and Narayanan, 2019) and granule (Beining et al., 2017b; Mishra and Narayanan, 2019; 2021c; b; Schneider et al., 2023) cells of the dentate gyrus, and stellate cells of the entorhinal cortex (Mittal and Narayanan, 2018; 2022). In addition, with reference to encoding functions of the hippocampal formation, degeneracy has been demonstrated in the emergence of short-(Mukunda and Narayanan, 2017) and long-term plasticity profiles in CA1 (Ashhad and Narayanan, 2013; Anirudhan and Narayanan, 2015) and DG (Shridhar et al., 2022), network decorrelation in the DG (Mishra and Narayanan, 2019; 2021c), place-cell tuning profiles in CA1 (Basak and Narayanan, 2018; 2020; Roy and Narayanan, 2021), spatial information transfer of CA1 place cells with rate (Roy and Narayanan, 2021) or phase (Seenivasan and Narayanan, 2020) coding, and in spatial coding functions of the dorso-ventral entorhinal axis (Pastoll et al., 2020).

In this study, our results extend previous lines of evidence for the manifestation of degeneracy in the emergence of characteristic physiological properties of DG granule cells (Beining et al., 2017b; Mishra and Narayanan, 2019; 2021c; b; Schneider et al., 2023). Our extension involved an extensive search of morphologically realistic granule cell models and covered a wide set of electrophysiological measurements that were all measured from the same set of biological granule cells (Mishra and Narayanan, 2020). This set of biological measurements allowed us to not just look at the distributions of individual measurements, but also ask if there were second- and higher-order relationships between model measurements and if they were comparable to experimental observations. We find the distributions and the co-dependencies of individual measurements to be comparable to their experimental counterparts (Figs. 2– 3), thus providing a valid population of granule models that match with their biological counterparts.

Our analyses demonstrate that these characteristic granule cell models emerge with their distributions spanning a wide range of parametric combinations with weak cross-dependencies in the parametric space (Figs. 5– 6). These observations translate to expansive degrees of freedom available to DG granule cells achieve characteristic physiological characteristics in spite of morphological and ion-channel distribution constraints. A fundamental advantage for the expression of degeneracy in biological system is the ability to achieve robust function through several disparate routes, thus reducing dependencies on any single component for executing precise function. These distinctions in the specific route taken to achieve characteristic functions might also translate to effective implementation of certain functions (Mishra and Narayanan, 2021b; c) or heterogeneities in other aspects (Shridhar et al., 2022) of granule cell physiology that account for the three-dimensional morphology.

### Future directions

The demonstration of degeneracy in morphologically realistic granule cell models, strongly constrained by several functional measurements and their co-dependencies, is a first step in understanding the physiology and pathophysiology of DG granule cells, their dendritic physiology, and network interactions. There are several specific directions that could be pursued with the understanding that the emergence of their physiology manifests degeneracy and that they form a complex system where functional specialization coexists with functional segregation. First, there are differences in the morphology, physiology, and the biophysical composition of DG granule cells in different locations and different states. These differences are known to exist along the dorso-ventral axis of the hippocampus (Desmond and Levy, 1982; 1985; van Groen et al., 2003; Schmidt et al., 2012; Strange et al., 2014; Lee et al., 2017; Schreurs et al., 2017; Cembrowski and Spruston, 2019; Botterill et al., 2021), pathological conditions (Wossink et al., 2001; Bender et al., 2003; Ellerkmann et al., 2003; Verina et al., 2007; Stegen et al., 2009; Young et al., 2009; Epsztein et al., 2010; Artinian et al., 2011; Stegen et al., 2012; Surges et al., 2012; Perederiy and Westbrook, 2013), and uniquely for DG granule cells, adult neurogenesis (Altman and Das, 1965; Doetsch and Hen, 2005; Zhao et al., 2008; Dieni et al., 2013; Aimone et al., 2014; Dieni et al., 2016; Beining et al., 2017a; Beining et al., 2017b; Li et al., 2017; Lodge and Bischofberger, 2019; Cole et al., 2020; Huckleberry and Shansky, 2021). The overall approach employed here could be used to build different heterogeneous populations of granule cells built with distinct morphologies and disparate sets of ion channels that emerge from respective experimental measurements that span a large set of characteristics from the same set of cells. Such analyses would enable a fundamental understanding of differences in heterogeneities, composition, extent of degeneracy in each state, and how they contribute to the physiology of the neurons and their networks (Ratté et al., 2014; Ratte and Prescott, 2016; Goaillard and Marder, 2021; Mishra and Narayanan, 2021d; Seenivasan and Narayanan, 2022; Marom and Marder, 2023; Mittag et al., 2023; Stober et al., 2023; Yang and Prescott, 2023).

Such insights about individual neuronal physiology and their composition could then be used to assess network scale functions such as different forms of decorrelation (Leutgeb et al., 2007; McHugh et al., 2007; Bakker et al., 2008; Clelland et al., 2009; Deng et al., 2010; Wick et al., 2010; Sahay et al., 2011; Schmidt et al., 2012; Vivar et al., 2012; Moreno-Bote et al., 2014; Wu et al., 2015; Cayco-Gajic et al., 2017; Chavlis et al., 2017; Mishra and Narayanan, 2019; 2021b; c) and engram cell formation (Liu et al., 2012; Ramirez et al., 2013; Redondo et al., 2014; Ramirez et al., 2015; Park et al., 2016; Roy et al., 2016; Titley et al., 2017; Lisman et al., 2018; Mishra and Narayanan, 2021d; 2022; Shridhar et al., 2022) assessed with morphologically realistic heterogeneous model populations of different neuronal subtypes in the hippocampus. The use of heterogeneous morphologically realistic models for all neuronal subtypes would provide deeper insights about degeneracy in the emergence of DG network function (Mishra and Narayanan, 2019; 2021c) and the role of active dendrites in mediating decorrelation as well as engram cell formation (Govindarajan et al., 2011; Chavlis et al., 2017). Finally, the use of such network models with morphologically realistic neurons that manifest degeneracy at different scales could be used to assess the multifarious and heterogeneous impact of different neuromodulators across different cells at different locations. In addition to changes mediated by neuromodulation, such morphologically realistic model populations could also be used to assess plasticity profile degeneracy (Anirudhan and Narayanan, 2015; Shridhar et al., 2022), plasticity heterogeneities (Shridhar et al., 2022), and plasticity degeneracy (Nagaraj and Narayanan, 2023) in implementing the encoding functions of the DG network.

## ACKNOWLEDGMENTS

The authors thank members of the cellular neurophysiology laboratory for helpful discussions and for comments on a draft of this manuscript. This work was supported by the Wellcome Trust-DBT India Alliance (Senior fellowship to R. N.; IA/S/16/2/502727), Department of Biotechnology (R. N.), and the Ministry of education (R. N. & S. K.).

## Author Contributions

S.K. and R.N. designed experiments; S.K. performed experiments; S.K. analyzed data; S.K. and R.N. wrote the paper.

## Competing Interest Statement

The authors declare that they have no competing interests.

## Notes

### Competing Interest Statement

The authors have declared no competing interest.

